# A Suite of Mouse Reagents for Studying Amelogenesis

**DOI:** 10.1101/2023.03.30.534992

**Authors:** Tomas Wald, Adya Verma, Victoria Cooley, Pauline Marangoni, Oscar Cazares, Amnon Sharir, Evelyn J. Sandoval, David Sung, Hadis Najibi, Tingsheng Yu Drennon, Jeffrey O. Bush, Derk Joester, Ophir D. Klein

## Abstract

Amelogenesis, the formation of dental enamel, is driven by specialized epithelial cells called ameloblasts, which undergo successive stages of differentiation. Ameloblasts secrete enamel matrix proteins (EMPs), proteases, calcium, and phosphate ions in a stage-specific manner to form mature tooth enamel. Developmental defects in tooth enamel are common in humans, and they can greatly impact the well-being of affected individuals. Our understanding of amelogenesis and developmental pathologies is rooted in past studies using epithelial Cre driver and knockout alleles. However, the available mouse models are limited, as most do not allow targeting different ameloblast sub-populations, and constitutive loss of EMPs often results in severe phenotype in the mineral, making it difficult to interpret defect mechanisms. Herein, we report on the design and verification of a toolkit of twelve mouse alleles that include ameloblast-stage specific Cre recombinases, fluorescent reporter alleles, and conditional flox alleles for the major EMPs. We show how these models may be used for applications such as sorting of live stage specific ameloblasts, whole mount imaging, and experiments with incisor explants. The full list of new alleles is available at https://dev.facebase.org/enamelatlas/mouse-models/.

## INTRODUCTION

Tooth enamel, the hardest tissue in the mammalian body, is one of the major biological innovations that has enabled radiation of mammals (Lacruz et al. 2017). Enamel has a highly hierarchical structure comprised of both crystalline and amorphous mineral components (DeRocher et al. 2020; Free et al. 2022). Amelogenesis is driven by dental epithelial cells called ameloblasts that are lost upon eruption of the tooth. Ameloblasts arise from dental stem cells (Harada et al. 1999; Sharir et al. 2019) and their terminal differentiation can be divided into four consecutive functionally distinct stages (Lacruz et al. 2017). In the presecretory stage, they change polarity and develop a protein-synthesizing apparatus. In the secretory stage, ameloblasts begin secreting enamel matrix proteins (EMPs), and inorganic ions (calcium and phosphate, among others). Thin mineral ribbons form in the secreted matrix, adjacent to the cell membrane as ameloblasts move away from the dentino-enamel-junction. Once the full thickness of the enamel layer has been delineated, the cells proceed via a brief transition stage to the maturation stage. Here the ameloblasts secrete Kallikrein 4 (KLK4) protease to degrade and remove EMPs (Hu et al. 2016; Lu et al. 2008). As the ameloblasts cycle through ruffle- and smooth-ended appearance, concomitant changes in enamel matrix pH and ion transport drive enamel crystallite growth in width and thickness (Robinson et al. 1995).

Finally, ameloblasts shrink and become the reduced enamel epithelium that adopts a protective function (Nanci 2012). Because amelogenesis is evolutionarily conserved, enamel formation in rodent models closely mirrors the process in humans. Unlike human teeth and rodent molars, in which differentiation and enamel secretion only occurs once before tooth eruption, rodent incisors grow continuously, due to the life-long presence of epithelial and mesenchymal progenitor cells (Krivanek et al. 2020; Sharir et al. 2019). The presence of all developmental stages along the length of the incisor makes it an ideal model for studying amelogenesis (**Figure S1A**) (Pugach and Gibson 2014).

Dissection of the mechanisms by which amelogenesis proceeds greatly benefits from tissue-specific Cre alleles. In prior work, epithelium specific *Krt5* and *Krt14* promoters (*Krt5*^*CreERT2*^, JAX:#029155; *Krt14*^*Cre*^, JAX:#018964; *Krt14*^*CreER*^, JAX:#005107) have often been used, however neither of them targets ameloblasts in a stage-specific manner. Cre driven recombination of conditional alleles affects all cell types in the enamel organ, precluding the ability to study ameloblasts specifically (**Figure S1B**). While some of these cell populations can be efficiently targeted using established Cre drivers (e.g. stratum intermedium by *Notch1*^*Cre*^, or *Krt17*^*Cre*^; actively-cycling progenitors by *Shh*^*Cre*^) (Seidel et al. 2010; Sharir et al. 2019), there is a lack of alleles targeting mature ameloblast subpopulations, reinforcing the need for tissue-specific Cre driver lines (Klein et al. 2017). To date, only three mouse models using the *Amelx* promoter have been published (Cho et al. 2013; Isono et al. 2021; Said et al. 2019).

Understanding the molecular mechanisms of amelogenesis in a tissue-specific, temporally resolved manner also requires targeting the main proteins secreted. Key EMPs are Amelogenin (AMEL) and Ameloblastin (AMBN) in the secretory stage; and Amelotin (AMTN) in the maturation stage (Bartlett et al. 2006; Fincham et al. 1999; Smith et al. 2017). AMEL and AMBN are proteolytically processed by matrix metalloproteinase 20 (MMP20) into multiple fragments to become the basis of the extracellular matrix (Iwata et al. 2007; Nagano et al. 2009). *Amelx, Ambn*, and *Mmp20* constitutive null mice displayed phenotypes with thin, hypomineralized or amorphous enamel layers lacking the rod/interrod microstructure (Hu et al. 2016; Smith et al. 2016; Wald et al. 2017). Ablation of *Amtn* or *Klk4* produces enamel that maintains a rod/interrod structure but is hypomineralized with severe defects in the surface structure (Hu et al. 2016; Nakayama et al. 2015). Thus, although constitutive knockout mouse models have been critical for establishing a foundational understanding of EMPs during amelogenesis, the almost total lack of mineral and severe defects make it difficult to infer the proteins’ detailed molecular roles. By contrast, conditional knockout mutants mediated by either epithelial or ameloblast stage-specific Cre drivers offer a path forward to precisely study the molecular mechanisms that drive amelogenesis.

Herein, we present the design, generation, and validation of 12 novel mouse alleles (**Figure S1C**). To allow targeting and manipulation of the different ameloblasts subpopulations, we generated five constitutive Cre and conditional CreERT2 alleles: *Ambn*^*Cre*^ and *Ambn*^*CreERT2*^, *Amelx*^*p2A-Cre*^, *Klk4*^*Cre*^, and *Klk4*^*CreERT2*^. To visualize and sort secretory and maturation ameloblasts, we have generated two fluorescent reporters: *Ambn*^*p2A-eGFP*^ (beginning in secretory stage) and *Klk4*^*mCherry*^ (maturation stage). Finally, to circumvent the severe phenotypes caused by constitutive knock out alleles, we have generated flox alleles for the major EMPs, *Ambn, Amelx*, and *Amtn*, and for the two major proteases, *Mmp20* and *Klk4*. These alleles will enable the study of a wide range of transcriptional, proteomic, and cell signaling events during amelogenesis at unprecedented resolution.

## RESULTS AND DISCUSSION

### Design and Verification of Stage-Specific Cre Alleles

We designed secretory stage-specific *Ambn*^*Cre*^, *Amelx*^*Cre*^, and *Ambn*^*CreERT2*^ and maturation stage-specific *Klk4*^*Cre*^ and *Klk4*^*CreERT2*^ alleles (**Figure 1**), using CRISPR/Cas9 technology. In heterozygous *Ambn* and *Klk4* null alleles, neither soft tissues nor enamel differ from wildtype (Hu et al. 2016; Wald et al. 2017) which allowed us to replace the 5’ portions of the structural genes for *Ambn* (**Figures 1A, I**) and *Klk4* (**Figures 1M, Q**) with the Cre or CreERT2 sequences followed by simian virus 40 late polyadenylation signal (SV40 polyA), a strong transcription terminator.

**Figure 1:**
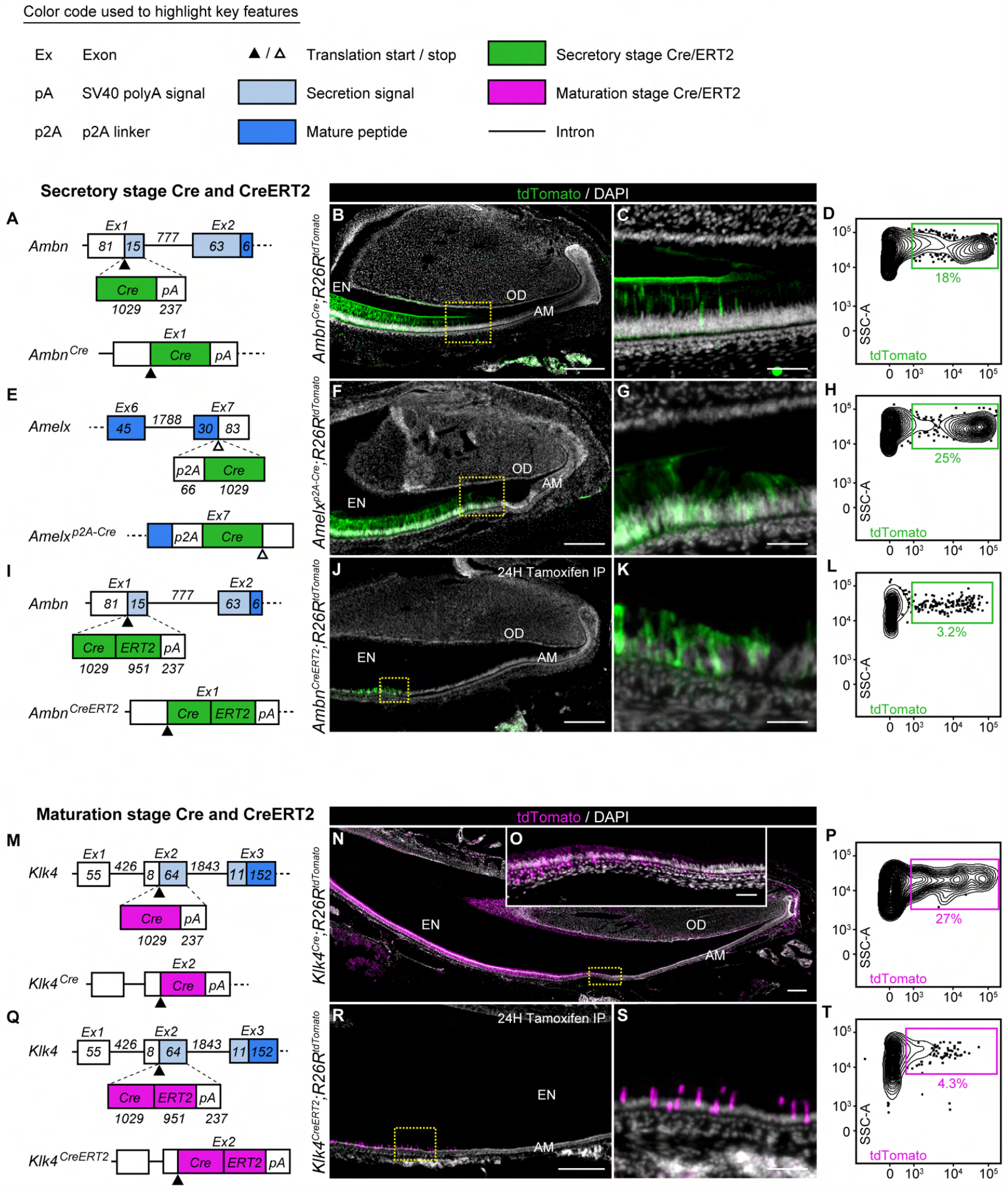
Design and verification of stage-specific constitutive and inducible Cre drivers. (**A, E, I, M, Q**) Schematic representation of the targeted loci, and design of the strategy to generate Cre alleles. (**B, F, J, N, R**) Composite fluorescence images of longitudinal sections of the mandibular incisors expressing constitutive Cre (**B, F, N**), and tamoxifen inducible CreERT2 (**J, R**) recombinase. Green pseudo-color corresponds to tdTomato expressed by secretory stage ameloblasts, magenta pseudo-color corresponds to tdTomato expressed by maturation stage ameloblasts. Nuclei were stained with DAPI (gray). Mice harboring CreERT2 were injected with tamoxifen 24 hours prior to tissue harvesting. OD: Odontoblasts; AM: Ameloblasts; EN: Enamel. Scale bar: 250 μm. (**C, G, K, O, S**) Higher magnification composite image of the region of interested highlighted by a rectangle in (**B, F, J, N, R**). Scale bar: 50 μm. (**D, H, L, P, T**) Flow cytometry analysis of tdTomato^+^ cells in incisor epithelium of mice expressing Cre (**D, H, P**), and CreERT2 (**L, O**) recombinase. tdTomato^+^ cells are indicated the colored box, and their fraction out of the total number of EpCAM^+^ cells is given. SSC-A: side scatter area.

*Amelx* resides on the X chromosome, rendering the KO/KI approach unsuitable. To leave the endogenous *Amelx* expression intact in females, we generated *Amelx*^*p2A-Cre*^, in which the p2A peptide causes ribosomes to skip during translation (**Figure 1E**) (Trichas et al. 2008). As a consequence, AMEL, with a C-terminal tag (the 19-mer peptide GSGATNFSLLKQAGDVEENPG), and Cre, are produced in a 1:1 ratio.

To verify the recombinase activity of the generated Cre alleles, mice were crossed with the *R26R*^*tdTomato*^ reporter strain (JAX:#007914) and mandibular sections were imaged using fluorescence microscopy. The tdTomato fluorescent protein is only expressed after elimination of a repressor by Cre mediated recombination, which allows analysis of the spatial distribution of Cre expression. In *Ambn*^*Cre/+*^;*R26R*^*tdTomato*^ and *Amelx*^*Cre/+*^;*R26R*^*tdTomato*^, fluorescence was first detected in a region where dentin and enamel start to be secreted, i.e. the secretory stage, and extended all the way through the reduced enamel epithelium (**Figures 1B-C and F-G**). In the incisor epithelium of *Klk4*^*Cre*^;*R26R*^*tdTomato*^, the onset of the tdTomato fluorescence was shifted in the incisal direction, to the region where the enamel reaches full thickness and ameloblasts transition into the maturation stage defined by secretion of KLK4 protease (**Figures 1N-O**) (Simmer et al. 2011). Interestingly, tdTomato was also detected in mesenchymal layer which is consistent with a previous report of *Klk4* expression in both epithelium and mesenchyme (Fukae et al. 2002).

To induce CreERT2 recombination, *Ambn*^*CreERT2*^ and *Klk4*^*CreERT2*^ mice were injected with one dose of tamoxifen 24 hours prior analysis (**Figures 1J, R**). The onset of tdTomato fluorescence in *Ambn*^*CreERT2*^;*R26R*^*tdTomato*^ incisor was shifted to greater distance from the cervical loop, and its expression was limited to a small patch of ameloblasts with a mosaic pattern (**Figures 1J-K**). In *Klk4*^*CreERT2*^;*R26R*^*tdTomato*^ the tdTomato recombination efficacy was also lower and occurred in a small subset of maturation-stage ameloblasts compared to its constitutive *Klk4*^*Cre*^ counterpart. We believe that the spatial shift of tdTomato expression to greater distance from the cervical loop observed in both CreERT2 mice reflects the time needed for nuclear accumulation of recombinase, and cellular accumulation of tdTomato protein.

Cre activity in all five models was further confirmed by flow cytometry analysis of TdTomato expression in epithelial (EPCAM^+^) cells. Inspection of histograms confirmed efficient recombination for the constitutive Cre drivers (**Figures 1D, H, P**). Consistent with the spatially restricted, likely mosaic expression observed by fluorescence microscopy, flow cytometry revealed a lower fraction of recombined cells in mice harboring CreERT2 variants (**Figure 1L and T**). While this demonstrated that inducible recombination is stage-specific, more work is required to establish how spatial expression and mosaicism depend on the tamoxifen dose and injection regimen.

### Design and Verification of Stage-Specific Fluorescent Reporter Alleles

While stage specific Cre driver lines are a powerful tool for functional studies, the resulting modification is permanent. Therefore, multiple transient cell populations such as secretory and maturation stage ameloblasts cannot be easily distinguished. We therefore generated fluorescent reporter alleles in which their expression is regulated by the promoter activity of endogenous genes specific to the secretory and maturation stage.

Specifically, we designed *Ambn*^*p2A-eGFP*^, and *Klk4*^*mCherry*^ reporter alleles. To leave the expression of endogenous *Ambn* intact, eGFP was fused to the 3’-end of *Ambn* via the p2A linker (**Figure 2A**). Because the p2A fusion approach was unsuccessful for *Klk4*, we replaced the structural portion of *Klk4* with *mCherry* followed by SV40 polyA (**Figure 2E**). Similarly, to the Cre alleles, neither of the fluorescent reporter alleles is expected to negatively impact the enamel layer when used as heterozygote.

**Figure 2:**
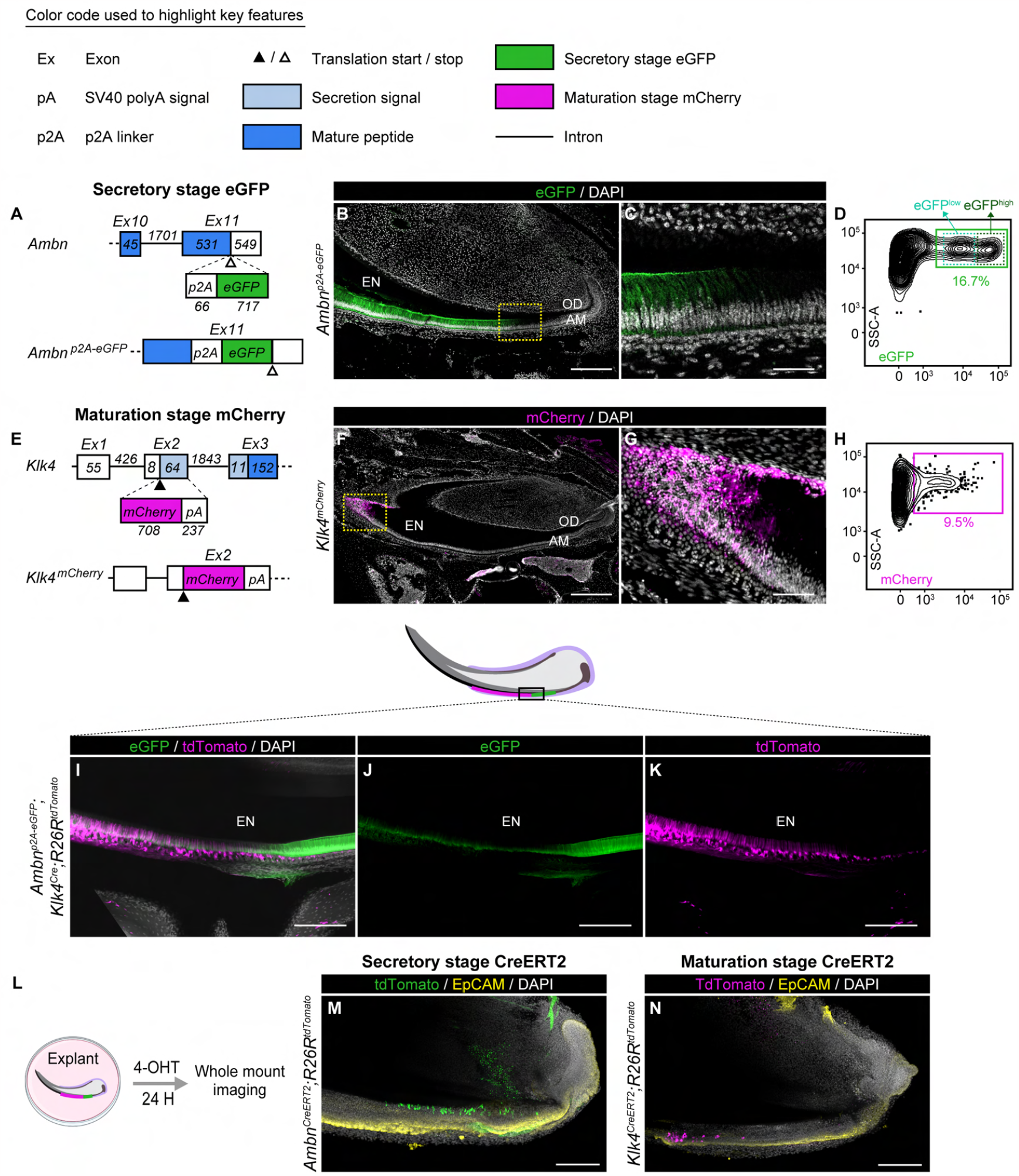
Design and verification of stage-specific fluorescent reporter alleles, and examples of the use of the novel alleles. (**A, E**) Schematic representation of the targeted loci, and design of the strategy to generate fluorescent reporter alleles. (**B, F**) Composite fluorescence images of longitudinal sections of the mandibular incisors expressing eGFP (green) (**B**); or mCherry (magenta) (**F**) fluorescent reporter. Nuclei were stained with DAPI (gray). OD: Odontoblasts; AM: Ameloblasts; EN: Enamel. Scale bar: 250 μm. (**C, G**) Higher magnification composite image of the region of interested highlighted by a rectangle in (**B, F**). Scale bar: 50 μm. (**D, H**) Flow cytometry analysis of eGFP^+^ (**D**), and mCherry^+^ (**H**) cells in incisor epithelium. eGFP^+^ and mCherry^+^ cells are highlighted by the rectangle, and their fraction out of the total number of EpCAM^+^ cells is given. SSC-A: side scatter area. (**I-K**) Whole mount multi-channel (**I**) and single-channel (**J-K**) fluorescence image of incisor of a mouse expressing *Ambn*^*p2A-eGFP*^;*Klk4*^*Cre*^;*R26R*^*tdTomato*^ alleles. Nuclei were stained with DAPI (gray). Scale bar: 200 μm. (**L**) Experimental schematics of whole mount imaging of an incisor explant established from lower hemimandible and grown *in vitro*. 24 hours following isolation, incisor was treated with 1 μM 4-OHT for 24 hours to induce Cre mediated recombination of *tdTomato* reporter gene. (**M-N**) Whole mount fluorescence images of incisors isolated from mouse expressing *Ambn*^*CreERT2*^;*R26R*^*tdTomato*^ (**M**), and *Klk4*^*CreERT2*^;*R26R*^*tdTomato*^ (**N**). Incisors were fixed and stained with anti-EpCAM antibody (yellow). Nuclei were counterstained with DAPI (gray). Scale bar = 200 μm.

Using fluorescence microscopy we verified the onset of eGFP expression in epithelial cells of *Ambn*^*p2A-eGFP*^ mouse approximately 500 μm incisal to the cervical loop (**Figure 2B-C**), where enamel matrix deposition begins. This closely matched the onset of the secretory stage as determined by immunohistochemical methods in the wildtype mouse (Wazen et al. 2009). Expression of eGFP was further confirmed by flow cytometry analysis, by which we identified almost 17% of EPCAM^+^ cells to be positive for eGFP (**Figure 2D**). Interestingly, we identified two ameloblast subpopulations that differed in eGFP intensity (eGFP^low^ and eGFP^high^), and therefore, expression levels of *Ambn*.

For the *Klk4*^*mCherry*^ mouse, we verified that the expression of mCherry is shifted incisally by 3000 μm, into the region where the enamel matrix has reached full thickness (**Figures 2F-G**). This matched onset of expression we observed in the *Klk4*^*Cre*^;*R26R*^*tdTomato*^ mouse. The fraction of epithelial (EPCAM^+^) cells that expressed mCherry in the *Klk4*^*mCherry*^ mouse using flow cytometry matched the high amount of the mCherry^+^ cells observed by fluorescence microscopy (**Figures 2H**).

### Applications: Identification and Isolation of Ameloblast Subpopulations

Identification of different ameloblast subpopulations in live tissues or for cell sorting is currently hampered by the lack of suitable surface epitopes that can be targeted by antibodies. We predicted that combination of stage-specific Cre and reporter alleles should allow us to overcome this challenge. We generated a “two-color” *Ambn*^*p2A-eGFP*^;*Klk4*^*Cre*^;*R26*^*tdTomato*^ mouse in which secretory stage ameloblasts were labeled by eGFP, while maturation stage ameloblasts and the reduced enamel epithelium were labeled by tdTomato. Inspection of sections by fluorescence microscopy confirmed the expression domains we observed previously, and also revealed the transition from secretory to maturation stage (**Figures 2I-K**). eGFP fluorescence tapered off quickly (∼200 μm) on the incisal side of the secretory stage but was still detectable at much reduced intensity at least 1 mm into the maturation stage (**Figure 2I**). This might have been caused either by continuous low expression of *Ambn* during the early maturation stage. Expression of tdTomato ramped up more gradually, over ∼500 μm, and its onset coincided well with the decay in eGFP (**Figure 2J**). This overlap of eGFP and tdTomato expression indicated a third, transitory stage ameloblasts.

Flow cytometry of EPCAM^+^ cells suggested there are five ameloblast subpopulations: eGFP^low^/tdTomato^-^, eGFP^high^/tdTomato^-^, eGFP^high^/tdTomato^low^, eGFP^low^/tdTomato^high^, and eGFP^-^/tdTomato^intermediate^ (**Figure S2A-B**). Expression of secretory stage-specific *Ambn*, and maturation stage-specific *Klk4* for some of these populations was evaluated by qPCR after FACS (**Figure S2C-D**). The first two populations likely represented the bulk of the secretory stage with *Ambn* strongly expressed; eGFP^low^ may have corresponded to early and eGFP^high^ to late secretory stage (**Figure S2B** green rectangles, **Figure S2C-D**). The comparatively small eGFP^high^/tdTomato^low^ population (orange rectangle) likely corresponded to transitory ameloblasts that arise from eGFP^high^ ameloblasts; with *Ambn* expression tailing off and that of *Klk4* increasing, the much larger population of eGFP^low^/tdTomato^high^ cells (magenta rectangle) was consistent with the fluorescence images of early maturation stage (**Figure S2B**). Expression of *Ambn* was similar to secretory stage and *Klk4* was strongly expressed (**Figure S2C-D**). Finally, as production of *Ambn* and *Klk4* is ceases (**Figure S2C-D**), the eGFP^-^/tdTomato^intermediate^ population (blue rectangle) likely corresponded to late maturation stage and the reduced enamel epithelium. In the future, combination of these alleles will enable detailed characterization of the different ameloblast subpopulations and their heterogeneity using the advantage of rapidly developing transcriptomic approaches.

The *Klk4*^*mCherry*^ allele allows detecting solely maturation stage-specific ameloblasts based on mCherry expression. Following isolation of mCherry^+^ cells, we observed high levels of *Klk4* expression in mCherry^+^ population compared to the rest of the epithelial cells (**Figure S2E**).

### Applications: Live Imaging of Incisor Epithelium in Explant Culture

Finally, we provide evidence that the alleles can be used for live imaging of ameloblast subpopulations. For instance, incisors dissected from the *Ambn*^*p2A-eGFP*^ mouse enabled time lapse imaging of secretory stage ameloblasts for up to 5 hours (**Figure S3, and videos S1-2**). In the future, this should allow us to test the common assumption that ameloblasts migrate collectively and that mineralization pattern preserves a complete record of their migratory paths (Cox 2010).

We envision that the alleles we described enable even more complex experiments. As an example, we established explant culture from incisors of *Ambn*^*CreERT2*^;*R26R*^*tdTomato*^ and *Klk4*^*CreERT2*^;*R26R*^*tdTomato*^ mice (**Figure 2L**), and induced recombinase activity by treating the explants with 4-hydroxytamoxifen 24 hours prior imaging (**Figures 2M-N**). The recombination of both *Ambn*^*CreERT2*^ and *Klk*^*CreERT2*^ alleles was mosaic similarly to what we observed *in vivo* (**Figures 1J, R**). We believe that the tdTomato signal in non-epithelial regions observed in the *Ambn*^*CreERT2*^;*R26R*^*tdTomato*^ incisor is coming from non-specific autofluorescence in mesenchymal region, as this hasn’t been observed *in vivo*.

### Design and Verification of Conditional Flox Alleles for Major EMPs and Proteases

To enable assessment of the respective molecular roles of genes forming enamel matrix, we designed flox alleles for major enamel matrix proteins *Ambn, Amelx*, and *Amtn*, and the two major proteases, *Mmp20* and *Klk4* (**Figure S1C**). As a result of common evolution they exhibit high degree of similarity at their 5’ ends and their exon-intron boundaries are in phase 0 (Kawasaki and Weiss 2003). To prevent exon skipping observed previously (Wazen et al. 2009) we targeted the exons containing either the translation start or secretion signal. To minimize the risk of disrupting the 5’ and 3’ splice sites, the LoxP sites were inserted approximately 100-200 base pairs (bp) up and downstream of the targeted exons (**Figures 3A-C**). Because *Amtn* exons 1 and 2 are separated only by a 105 bp intron, we inserted the 5’ LoxP site 355 bp upstream of exon 1, and the 3’ LoxP site after exon 2 (**Figure 3C**).

**Figure 3:**
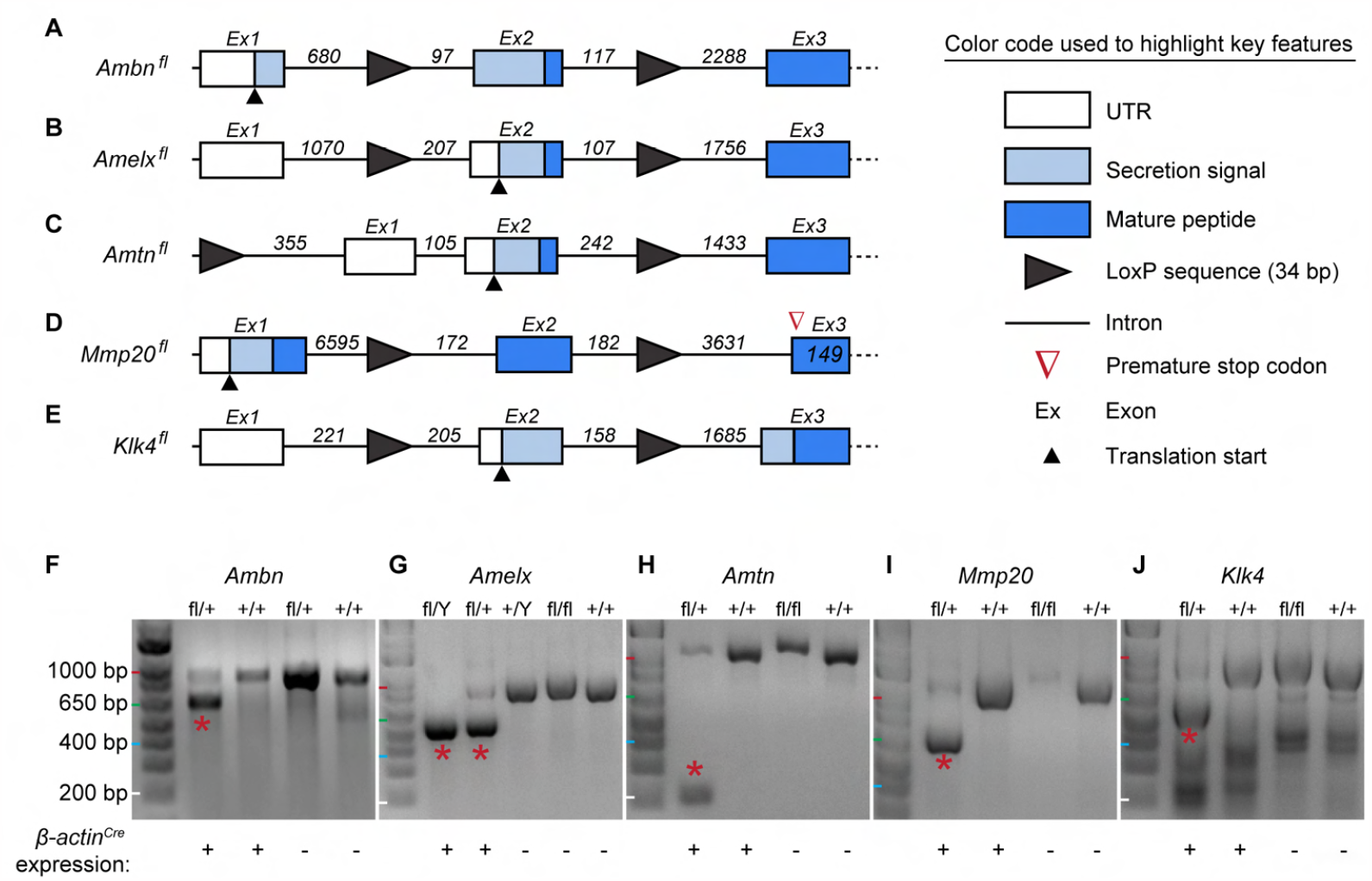
Design and verification of conditional alleles. (**A-E**) Schematic representation of the structure of flox alleles for the major enamel matrix genes and proteinases. (**F-J**) PCR based verification of LoxP recombination using the constitutively active and ubiquitously expressed β*-actin*^*Cre*^ driver. β*-actin*^*Cre*^-mediated recombination of LoxP sites results in excision of the sequence between LoxP sites which results in shortening of the amplified PCR product corresponding to null allele (red asterisks). fl/+: LoxP heterozygous; fl/fl: LoxP homozygous; fl/Y: LoxP hemizygous; +/+: Control; Cre^+^: β*-actin*^*Cre*^ allele. Cre^-^: no presence of β*-actin*^*Cre*^ allele.

*Mmp20* is localized on mouse chromosome 9 and *Klk4* on chromosome 7. The genomic structure of *Mmp20* allowed inserting LoxP sites up and downstream of exon 2 which leads to insertion of a premature stop codon in exon 3. The resulting aberrant MMP20 translated protein is composed of only 22 amino acids (**Figure 3D**). KLK4 is a small protease consisting of 222 amino acids encoded by 6 exons. Here, the LoxP sites were inserted up and downstream of exon 2, which encodes the translation start and most of the secretion signal (**Figure 3E**).

To verify the successful recombination of the LoxP sites, F1 mice were crossed with mice harboring the ubiquitously expressed β*-actin*^*Cre*^ (JAX, #019099). β*-actin*^*Cre*^ allows testing the LoxP functionality by PCR analysis of tail snip lysates of mice harboring flox alleles by generating two PCR products: the longer one corresponded to the wildtype allele, and the shorter one corresponded to the null allele (**Figures 3F-J**). In contrast, PCR analysis of littermates negative for either β*-actin*^*Cre*^ or floxed gene did not produce the shorter product. We concluded that all the new flox alleles can be used for functional studies when crossed with appropriate Cre drivers.

### Applications: Conditional deletion of enamel matrix genes in enamel epithelium

Three-dimensional reconstructions of hemimandibles generated by synchrotron micro-computed tomography (SMCT) revealed no differences in gross tooth morphology between wildtype and mice homozygous for floxed alleles (**Figures 4A, C, E, G, I**). Incisors from unrecombined flox mice displayed an increase in enamel thickness in the secretory stage followed by an increase in mineral density in the maturation stage, closely resembling the control hemimandible. Scanning electron microscopy (SEM) of etched incisor cross sections showed that the decussating rod/interrod structure of inner enamel was present in the mutants (**Figures 4B, D, F, H, J**), and that the outer enamel had a structure matching the wildtype (not shown). Together, these data showed that normal enamel production is not disrupted by the insertion of the LoxP sites.

**Figure 4:**
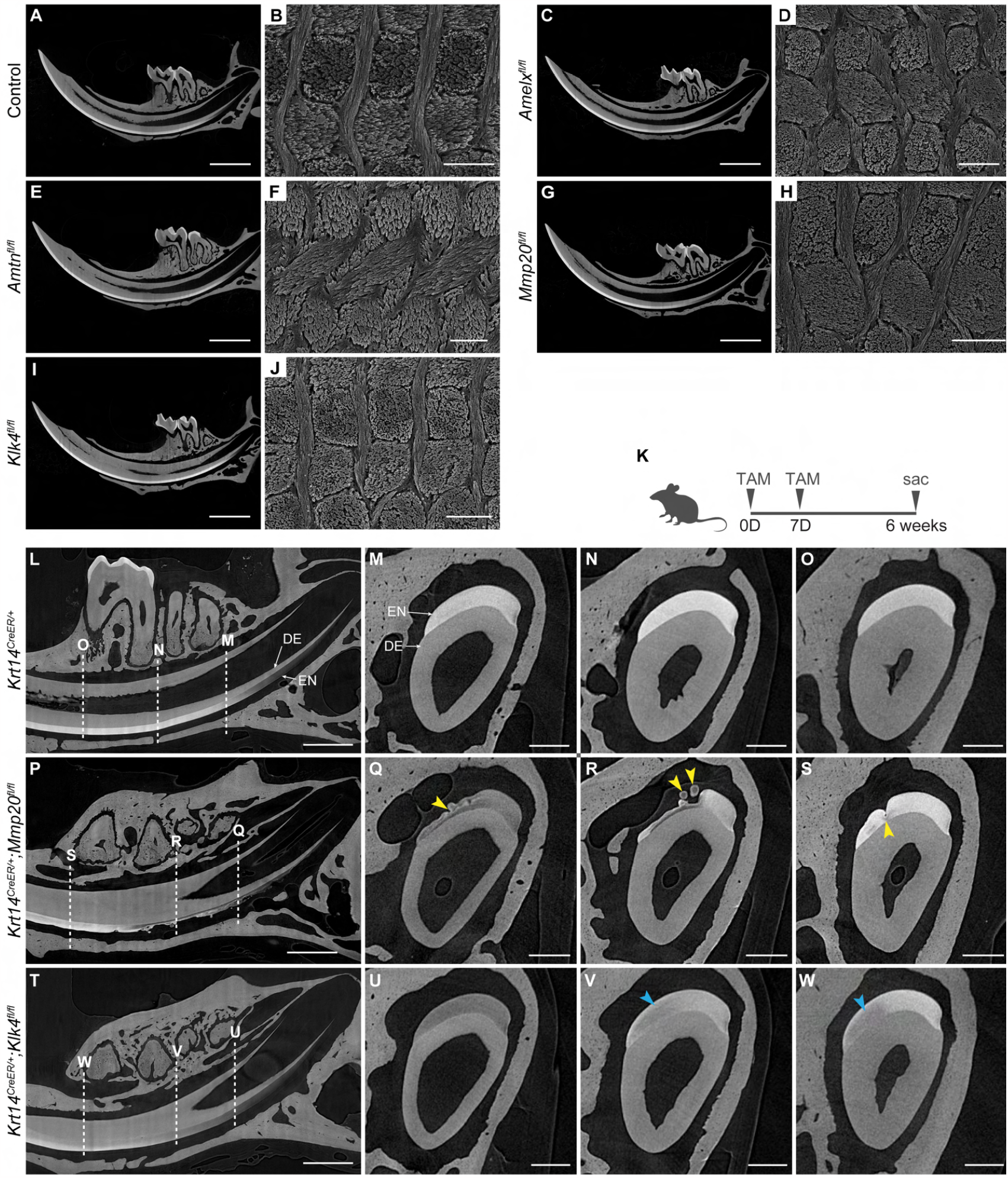
Insertion of LoxP sequences has no negative effect on the enamel microstructure by itself, but Cre mediated genetic deletion of floxed alleles results in severely impaired enamel layer. (**A, C, E, G, I**) Virtual sections, approximately parallel to the sagittal plane, through SMCT reconstruction of whole hemimandibles from (**A**) control, (**C**) *Amelx*^*fl/fl*^, (**E**) *Amtn*^*fl/fl*^, (**G**) *Mmp20*^*fl/fl*^, and (**I**) *Klk4*^*fl/fl*^ mice. Scale bar = 1 mm. (**B, D, F, H, J**) Scanning electron microscopy images of ground and polished, etched incisor enamel sections from (**B**) control, (**D**) *Amelx*^*fl/fl*^, (**F**) *Amtn*^*fl/fl*^, (**H**) *Mmp20*^*fl/fl*^, and (**J**) *Klk4*^*fl/fl*^ mice. Scale bar = 2 μm. (**K**) Schematic detailing tamoxifen (TAM) injection timepoints before sacrifice (sac). (**L, P, T**) Virtual sections, approximately parallel to the sagittal plane, through SMCT reconstruction of whole hemimandibles from (**L**) *Krt14*^*CreER/+*^, (**P**) *Krt14*^*CreEr/+*^*;Mmp20*^*fl/fl*^, and (**T**) *Krt14*^*CreER/+*^*;Klk4* ^*fl/fl*^ mice. Incisor dentin (DE) and enamel (EN) are labeled. Scale bar = 2 mm. (**M-O, Q-S, U-W**) Virtual sections, approximately parallel to the coronal plane (normal to the long axis of the incisor) at locations indicated by dashed lines in (**L**) for *Krt14*^*CreER/+*^ (**M-O**), *Krt14*^*CreEr/+*^*;Mmp20*^*fl/fl*^ (**Q-S**), and *Krt14*^*CreER/+*^*;Klk4*^*fl/fl*^ (**U-W**) mice. Incisor dentin (DE) and enamel (EN) are labeled. Yellow arrowheads represent grooves or ectopic mineral growths, and blue arrowheads represent hypomineralized regions. Scale bars = 250 μm.

Finally, we sought to demonstrate the ability to use tamoxifen to induce recombination targeting the two key proteases in a subset of ameloblasts. For this purpose, we generated *Krt14*^*CreER*^;*Mmp20*^*fl/fl*^ and *Krt14*^*CreER*^;*Klk4*^*fl/fl*^ mice, which were injected with two doses of tamoxifen (**Figure 4K**) and analyzed 7 weeks later (**Figures L-W**). After this time, we expect that the incisor and the secretory and maturation stage enamel epithelium have completely turned over, meaning that the phenotype observed is due to an ameloblast layer that lacks the corresponding protease.

The expected phenotype for constitutive *Mmp20* null mice is that of a thin, heterogeneous incisor enamel layer with an irregular, bumpy surface, a band of severe hypomineralization close to the DEJ, and surface nodules with higher mineral density (Hu et al. 2016). In the conditional *Mmp20* null mutants, regions that show exactly this phenotype can be observed in close proximity (the same transverse slice) to regions in which enamel seems to have grown to full thickness, and with a mineral density close to the expected (**Figures 4P, Q**). Interestingly, in other areas enamel defects included deep grooves and a locally hypomineralized layer at the DEJ even though enamel thickness and mineral density seemed mostly normal. This phenotype might have been caused by expression of *Mmp20* in mesenchyme, as it was published earlier (Sharir et al. 2019). The expected phenotype for *Klk4* constitutive null mice is full thickness enamel with a significantly reduced mineral density throughout (Hu et al. 2016). Consistent with mosaic loss of *Klk4* expression after tamoxifen induction, we observed stage-appropriate enamel thickness and mineralization in secretory stage (**Figure 4U**). However, in some regions, mineralization does not increase in maturation stage, resulting in full thickness enamel with hypomineralized patches (**Figures 4V, W**). Note that because of the stochastic nature of tamoxifen recombination in a continuously regenerating tissue, the phenotype is variable when comparing between teeth in the same mouse, or between different animals. Several options for fine tuning exist, including tamoxifen dosage, selection of Cre driver, or choosing the time delay between dosing and imaging. We envision that this may ultimately enable experiments in which specific functions are ablated for a small cluster of ameloblasts or even a single cell; this may help understand aspects such as formation of enamel rods and the collective ameloblast migration that gives rise to the decussating pattern of inner enamel.

To conclude, we report generation of novel ameloblasts stage-specific alleles. We envision that these alleles will be used for characterization of molecular mechanisms involved in ameloblast biology, and in hydroxylapatite deposition. Moreover, their combination with various flox alleles can be used to ablate genes that are involved in dental malformations in a spatially and temporally restricted manner with little to no off-target effects. The full list of ameloblast stage specific alleles is available at https://dev.facebase.org/enamelatlas/mouse-models/, and new models will be added here when available.

## ACKNOWLEDGEMENTS

We thank Brooks Hoehn, Asoka Rathnayake, Aimee Cortez, and Sergio Lopez for technical assistance. We would like to acknowledge the Transgenic Gene Targeting Core at Gladstone Institutes and the core director Junli Zhang for generating the mutant alleles, and the staff at the Biological Imaging Development CoLab (BIDC) at UCSF for their support. We acknowledge the PFCC (RRID:SCR_018206) supported in part by Grant NIH P30 DK063720 and by the NIH S10 Instrumentation Grant S10 1S10OD021822-01. This work was funded by NIH UG3DE028872 to O.D.K. and D.J. This work was in part supported by an NSF GRFP fellowship to V.C. (DGE-1842165). Part of this work was performed at the DuPont-Northwestern-Dow Collaborative Access Team (DND-CAT) located at Sector 5 of the Advanced Photon Source (APS). DND-CAT is supported by Northwestern University, The Dow Chemical Company, and DuPont de Nemours, Inc. This research used resources of the Advanced Photon Source, a U.S. Department of Energy (DOE) Office of Science User Facility operated for the DOE of Science by Argonne National Laboratory under Contract No. DE-AC02-06CH11357. This work made use of the following core facilities operated by Northwestern University: the EPIC facility of Northwestern University’s NUANCE Center, which have received support from the SHyNE Resource (NSF ECCS-2025633), the IIN, and Northwestern’s MRSEC program (NSF DMR-1720139); and MatCI, supported by the MRSEC program (NSF DMR-1720139) at the Materials Research Center.

## AUTHOR CONTRIBUTIONS

Tomas Wald: Contributed to conception, design, data acquisition and interpretation, performed all statistical analyses, drafted and critically revised the manuscript. Adya Verma: Contributed to design, data acquisition and interpretation, drafted and critically revised the manuscript. Victoria Cooley: Contributed to design, data acquisition and interpretation, drafted and critically revised the manuscript. Pauline Marangoni: Contributed to conception, data acquisition and interpretation, and critically revised the manuscript. Oscar Cazares: Contributed to data acquisition and interpretation, and critically revised the manuscript. Amnon Sharir: Contributed to conception, and critically revised the manuscript. Evelyn J. Sandoval: Contributed to data acquisition, interpretation, and critically revised the manuscript. David Sung: Contributed to data acquisition, interpretation, and critically revised the manuscript. Hadis Najibi: Contributed to data acquisition and critically revised the manuscript. Tingsheng Yu Drennon: Contributed to data acquisition and critically revised the manuscript. Jeffrey O. Bush: Contributed to conception, design, and critically revised the manuscript. Derk Joester: Contributed to conception, design, and critically revised the manuscript. Ophir D. Klein: Contributed to conception, design, and critically revised the manuscript. All authors gave their final approval and agreed to be accountable for all aspects of the work.

## METHODS

Methods, and genotyping protocols are listed in the supplemental material. We attest that we have complied with the Reporting In Vivo Experiments (ARRIVE) 2.0 checklist.

