## Supplemental material for "A Suite of Mouse Reagents for Studying Amelogenesis"

**Figure S1: All stages of enamel development can be observed along the cervical-incisal axis of the continuously growing mouse incisor.**

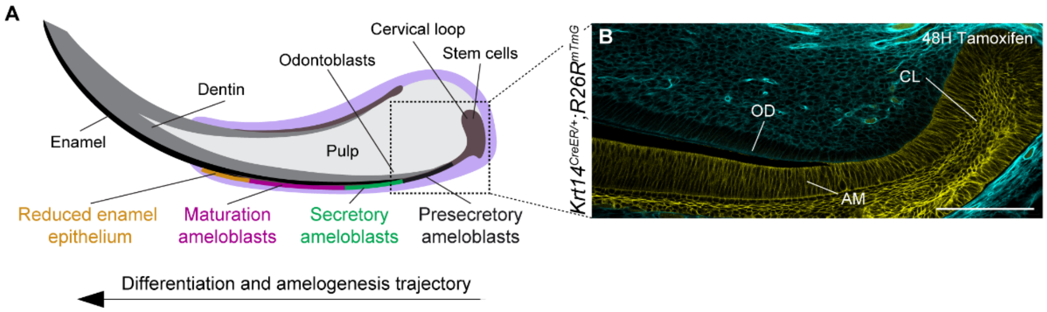


**C**

LIST OF GENERATED ALLELES SPECIFIC FOR SECRETORY AND MATURATION STAGE AMELOBLASTS, AND THEIR CHROMOSOMAL LOCALIZATION

| Cre drivers | | Fluorescent reporters | | FLOX | |
|---|---|---|---|---|---|
| *Amelx*$^{p2A\text{-}Cre}$ | (Ch. X) | *Ambn*$^{p2A\text{-}eGFP}$ | (Ch. 5) | *Amelx*$^{fl}$ | (Ch. X) |
| *Ambn*$^{Cre}$ | (Ch. 5) | *Klk4*$^{mCherry}$ | (Ch. 9) | *Ambn*$^{fl}$ | (Ch. 5) |
| *Ambn*$^{CreERT2}$ | (Ch. 5) | | | *Mmp20*$^{fl}$ | (Ch. 7) |
| *Klk4*$^{Cre}$ | (Ch. 9) | | | *Amtn*$^{fl}$ | (Ch. 5) |
| *Klk4*$^{CreERT2}$ | (Ch. 9) | | | *Klk4*$^{fl}$ | (Ch. 9) |

**Figure S1: All stages of enamel development can be observed along the cervical-incisal axis of the continuously growing mouse incisor.** (**A**) Schematic drawing of the mandibular mouse incisor. Dental epithelial stem cells located in the labial cervical loop (CL) differentiate and give rise to secretory stage ameloblasts. These secrete enamel matrix proteins and proteases which are responsible for hydroxylapatite deposition. Once full thickness of enamel is reached, secretory stage ameloblasts undergo differentiation into maturation stage ameloblasts to complete the mineralization of the hydroxylapatite. Finally, maturation stage ameloblasts differentiate into reduced enamel epithelium. Amelogenin (AMEL), Ameloblastin (AMBN), Enamelin (ENAM), and Matrix metalloprotease 20 (MMP20) are the key proteins secreted during the secretory stage. Structural protein Amelotin (AMTN) and protease Kallikrein 4 (KLK4) are secreted during the maturation stage. (**B**) Immunofluorescence image of a longitudinal section of a mandibular incisor of a *Krt14*$^{CreER/+}$;*R26R*$^{mTmG}$ mouse injected with tamoxifen 48 hours prior to tissue harvesting. Recombination of *R26R*$^{mTmG}$ occurs in the entire incisor epithelium. Scale bar = 200 µm. CL: Cervical loop; OD: Odontoblasts; AM: Ameloblasts. (**C**) List of mutant alleles generated and described in the current study and their chromosomal localization. Green-colored alleles are relevant to secretory stage ameloblasts. Magenta-colored alleles are relevant to maturation stage ameloblasts.

**Figure S2: Analysis of *Ambn* and *Klk4* transcripts by qPCR in the respective ameloblast subpopulations isolated from *Klk4*$^{Cre/+}$;*Ambn*$^{eGFP/+}$;*R26R*$^{tdTomato}$ and *Klk4*$^{mCherry/+}$ mice.**

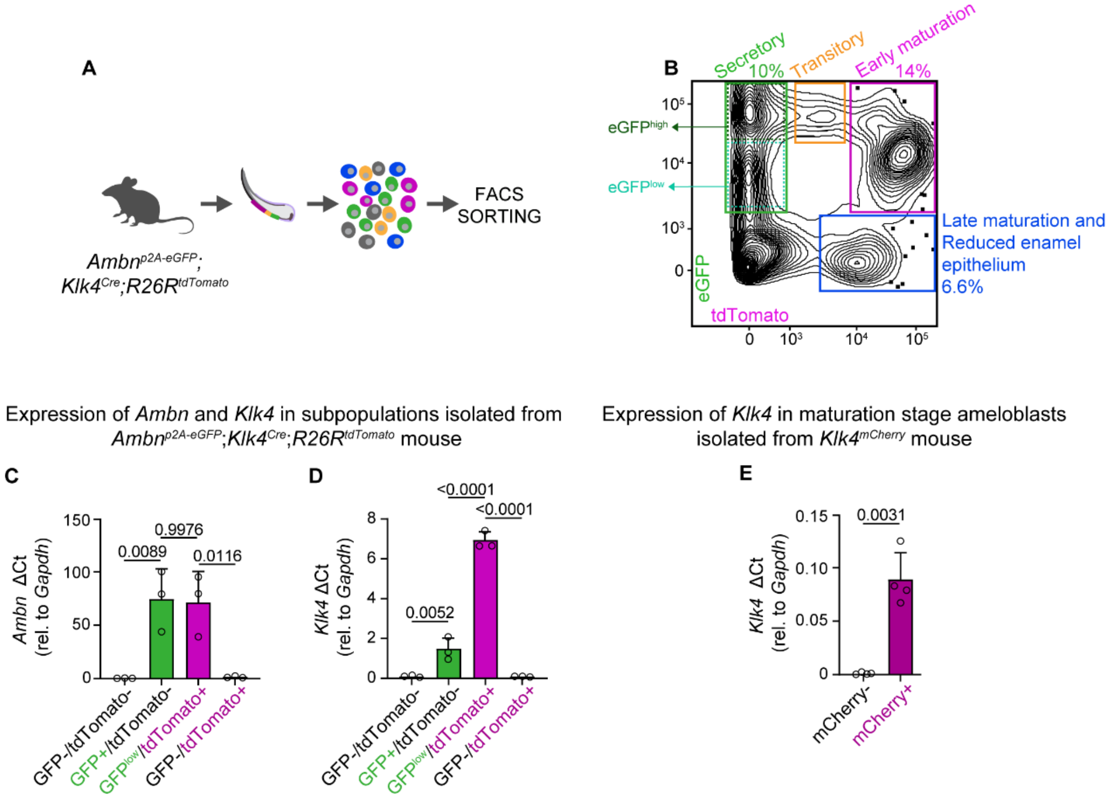


**Figure S2: Analysis of *Ambn* and *Klk4* transcripts by qPCR in the respective ameloblast subpopulations isolated from *Ambn*$^{eGFP/+}$;*Klk4*$^{Cre/+}$;*R26R*$^{tdTomato}$ and *Klk4*$^{mCherry/+}$ mice.** (**A**) Experimental schematics for FACS-sorting of secretory, transitory and maturation stage ameloblasts from mouse incisor using *Ambn*$^{p2A-eGFP}$;*Klk4*$^{Cre}$;*R26R*$^{tdTomato}$ alleles. (**B**) 2D histogram of ameloblasts subpopulations distinguished by the expression of eGFP and tdTomato. Secretory (eGFP$^{low}$/tdTomato$^{-}$ and eGFP$^{high}$/tdTomato$^{-}$), transitory (eGFP$^{high}$/tdTomato$^{+}$), early maturation stage (eGFP$^{low}$/tdTomato$^{+}$), and late maturation stage and reduced enamel epithelial cells (eGFP$^{-}$/tdTomato$^{+}$) are delineated, and their fraction of the total number of epithelial EpCAM$^{+}$ cells is given. (**C-E**) Bar plots of the difference of the qPCR cycle threshold (ΔCt) for *Ambn* (**C**), and *Klk4* (**D-E**) compared to that of Glyceraldehyde 3-phosphate dehydrogenase (*Gapdh*). Live EpCAM$^{+}$ cells were isolated from incisors of *Ambn*$^{eGFP/+}$;*Klk4*$^{Cre/+}$;*R26R*$^{tdTomato}$ mouse (**C-D**) into four groups based on the presence of GFP and/or tdTomato expression; cells isolated from *Klk4*$^{mCherry/+}$ mouse incisors (**E**) were sorted into two groups based on the presence of mCherry. Open circles = biological replicates. Error bars = standard deviation. N = 3-4 mice per condition.

**Figure S3: Live imaging of the mouse incisor expressing *Ambn*$^{p2A\text{-}eGFP}$.**

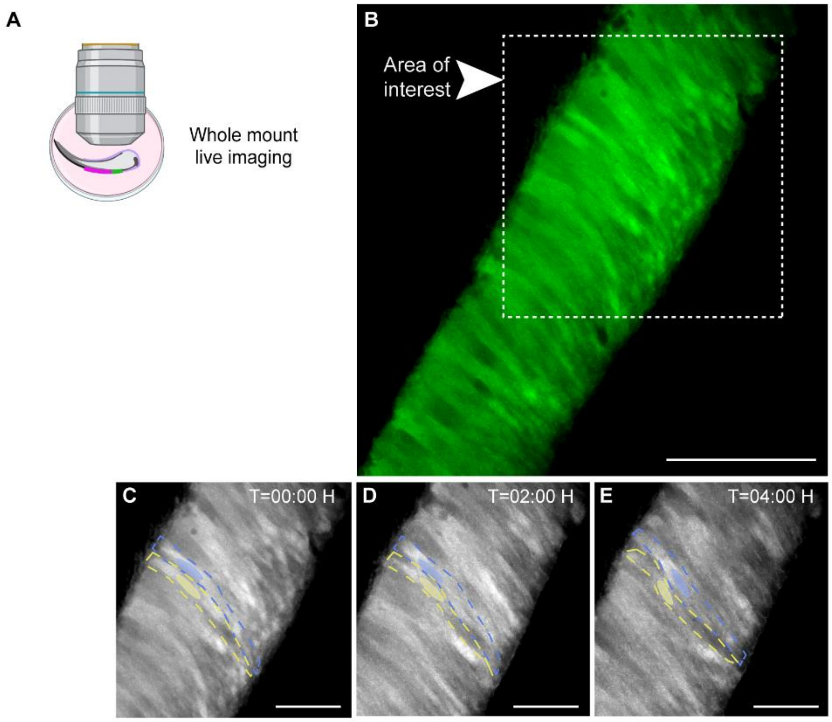


**Figure S3: Live imaging of mouse incisor expressing *Ambn*$^{p2A\text{-}eGFP}$.** (**A**) Experimental schematics for whole mount live imaging of an incisor from *Ambn*$^{p2A\text{-}eGFP}$ expressing mouse. (**B**) Fluorescence image of live whole mount at $t$ = 0 of video S1. The area indicated represents the field of view of the time lapse sequence shown in Figures S3C-E and video S2. Scale bar = 50 µm. (**C-E**) Snapshots of wholemount live imaging at time 0 hours (**C**), 2 hours (**D**), and 4 hours (**E**). Tracked cells are highlighted by blue- and yellow-colored ovals. Regions with actively moving blue- and yellow highlighted cells is marked by the dotted line. Images relate to videos S1-2. Scale bar: 25 µm.

**Video S1, related to Figure S3:** Timelapse movie of live ameloblasts in whole mount of *Ambn*$^{eGFP/+}$ incisor, recorded by two-photon microscopy. Green pseudo-color corresponds to eGFP expressed by secretory stage ameloblasts.

**Video S2, related to Figure S3:** Timelapse movie of live ameloblasts in whole mount of *Ambn*$^{eGFP/+}$ incisor, recorded by two-photon microscopy. Higher magnification (200%) video in pseudo-colors of the original Video S1 showing movement of ameloblasts from the apical to basal position. The magnified region corresponds to the area highlighted in Figures S3C-E.

### METHODS

#### Animals

All experimental procedures involving mice were carried out in accordance with approved protocols by the Institutional Animal Care and Use Committee (IACUC) and Laboratory Animal Resource Center (LARC) at the University of California San Francisco. Mice were handled in accordance with the principles and procedures of the Guide for the Care and Use of Laboratory Animals under the approved protocol AN195665. All mouse strains were maintained on a C57BL6/J background. Animals were housed under pathogen-free conditions in 12-hour light/dark cycles at 23 ± 1°C and humidity 55 ± 15%. Food and water were provided *ad libitum*. Animals were weaned 21 to 28 days after birth, handled and euthanized according to approved procedures. Mice were at least 8 weeks old at the time of experiments and cell isolations, which is considered the standard adult age in incisor biology (Sharir et al., 2019).

#### Genome editing

Genome edited mouse models were generated in collaboration with the Transgenic Gene Targeting Core at Gladstone Institutes, CA, on a C57BL/6 background (Charles River Laboratories) by microinjection or electroporation of 1-day old zygotes with Ribonucleoprotein complex (composed of Cas9, tracrRNA and one or two crRNAs) together with single-stranded DNA. The crRNAs of highest score and specificity were designed using Alt-R CRISPR-Cas9 design tools (https://www.idtdna.com/site/order/designtool/index/CRISPR_PREDESIGN). Genomic sequences were downloaded from ENSEMBL.org (genome assembly GRCm39): ***Ambn***: ENSMUSG00000029288; ***Amelx***: ENSMUSG00000031354; ***Amtn***: ENSMUSG00000029282; ***Mmp20***: ENSMUSG00000018620; ***Klk4***: ENSMUSG00000006948. crRNA and single stranded DNA was designed in Snapgene. The following 23-bp sequences including PAM recognition sites, which are underlined in 5'-3' orientation were used to produce crRNA: $\boldsymbol{Ambn^{Cre}}$ and $\boldsymbol{Ambn^{CreERT2}}$: ACAGTGAATGTCAGCATCTAAGG; $\boldsymbol{Amelx^{p2A\text{-}Cre}}$: CTTATCTTAACAGCTTAACATGG and TGGGAAGGTAGGAGAGTGGTTGG; $\boldsymbol{Klk4^{Cre}}$, $\boldsymbol{Klk4^{CreERT2}}$ and $\boldsymbol{Klk4^{mCherry}}$: TCGTGCAGTGACCATCATGTTGG and TGCCTCATCCTTGAGGTCACAGG; $\boldsymbol{Ambn^{p2A\text{-}}}$

$^{eGFP}$: CTGCTATGTCAAGGTTATCAGGG and TTTTAGCAACAAGGTACTATGGG; **$Ambn^{flox}$**: CTGTTCTTCCAAAGACAATCAGG and TGTAGCTCCCAGTGAGTCAATGG; **$Amelx^{flox}$**: GCTGAAGGTTAACGAGTTAGAGG and TATATCCCCAAGTATCTAATAGG; **$Amtn^{flox}$**: TCAAGGAAACAAGTAAGGTTTGG and AAAGCAATATAGCAGCCCTGAGG; **$Mmp20^{flox}$**: ACAAGTCTTAGTACTACTCAAGG and CCATTACAAGTCTGACCAGTTGG; **$Klk4^{flox}$**: GGAGGCATAGATCTTCACTTAGG and AGTCTTAGTGGTACCACTAAGGG. The crRNAs and tracrRNA (IDT, #1072532) were mixed in equimolar ratio and assembled into a ribonucleoprotein (RNP) complex with Cas9 protein (IDT; #1081058), microinjected or electroporated into 1-cell zygotes together with single stranded DNA (synthesized in IDT or Genewiz) and transferred into pseudopregnant foster mice. Putative founders were analyzed by PCR and sequencing. The crRNA off-target sites were identified using http://chopchop.cbu.uib.no/ and those encoded on the targeted chromosome were sequenced. The positively targeted founder F0 mice (usually between 1 and 3 founders) were backcrossed with C57BL6/J mice to generate stable F1 and F2 progenies. The F2 progeny were used for allele verification and sperm cryopreservation.

**Genotyping**

Approximately 1 mm of tail from 13-17 days old mice was snipped with razor blade and lysed for 12-16 hours at 60°C in 100 µl of lysis buffer (VIAGEN, #102-T) containing 0.5 µl of Proteinase K (Sigma, #P6556-500MG, stock concentration 20 mg/mL. Lysates were diluted with 450 µl d$H_2O$ and stored in 4°C. Genotyping was done by routine PCR reaction by using 2X GoTaq polymerase mix (Promega, #M7122). PCR reaction was assembled as follows: 15 µl d$H_2O$, 15 µl 2X GoTaq, 0.5 µl of forward and reverse primers (100 µM), and 2.5 µl tail lysate. PCR program used for genotyping all alleles: 1) Initial denaturation at 94°C for 90 sec; 2) 35 cycles of PCR elongation: Denaturation at 94°C for 30 sec; Annealing at 58-62°C for 30 sec; Elongation at 72°C for 35-60 sec. 3) Final elongation at 72°C for 60 sec. Primers used for PCR genotyping in 5'-3' orientation: **$Ambn^{Cre}$**: Forward-WT: GTCAGCATCTAAGGTAAAATG; Forward-Cre: GATTGATTTACGGCGCTAAG; Reverse-common: CTGATCAGCAGGTGTGCTTATC. PCR product size: WT: 279 bp, Cre: 725 bp. WT and Cre alleles can be analyzed in single PCR reaction. **$Ambn^{CreERT2}$**: Forward-WT: GTCAGCATCTAAGGTAAAATG; Forward-CreERT2:

CTGATCAGCAGGTGTGCTTATC; Reverse-common: GGCTCTACTTCATCGCATTC. PCR product size: WT: 269 bp, CreERT2: 576 bp. WT and CreERT2 alleles must be analyzed in separate PCR reactions. ***Amelx**$^{p2A\text{-}Cre}$*: Forward-WT: CAACAGAAAAATGATTCCAACC; Forward-Cre: CGATTGATTTACGGCGCTAAGG; Reverse-common: AACCCTTATTGATAAGGTTACAG. PCR product size: WT: 235 bp, Cre: 446 bp. WT and Cre alleles can be analyzed in single PCR reaction. ***Klk4**$^{Cre}$*: Forward-WT: GTGCCTCATCCTTGAGGTC; Forward-Cre: GATAGTGAAACAGGGGCAAT; Reverse-common: GAATCAACTAGTCACTCAGG. PCR product size: WT: 247 bp, Cre: 507 bp. WT and Cre alleles can be analyzed in single PCR reaction. ***Klk4**$^{CreERT2}$*: Forward-WT: GTGCCTCATCCTTGAGGTC; Forward-CreERT2: GGCTCTACTTCATCGCATTC; Reverse-common: GAATCAACTAGTCACTCAGG. PCR product size: WT: 247 bp, CreERT2: 534 bp. WT and CreERT2 alleles must be analyzed in separate PCR reactions. ***Ambn**$^{p2A\text{-}eGFP}$*: Forward-WT: CAAGAGCCCTGATAACCTTGA; Forward-eGFP: CAAGAGCCGGGAAGCGGAGC; Reverse-common: ACAGCTACATGCTATACCACC. PCR product size: WT: 291 bp, eGFP: 1074 bp. WT and eGFP alleles can be analyzed in single PCR reaction. ***Klk4**$^{mCherry}$*: Forward-mCherry: CTAAGTGAAGATCTATGCCTC; Reverse-mCherry: CATGGTCTTCTTCTGCATTAC. PCR product size: mCherry: 656 bp. ***Ambn**$^{flox}$*: Catch-all PCR: Forward: GCACACCTGCTGATCAGATC; Reverse: CTAGCAAGGACTTGATAATAAAAC. PCR product size: WT: 901 bp, flox: 969 bp, Null allele: 651 bp. Flox PCR: Forward: GCACACCTGCTGATCAGATC; Reverse: CTTCCAAAGACAATAACTTCG. Flox PCR product size: 477 bp. WT PCR: Forward: CCCAACTGTGTTTTAGATTCCA; Reverse: GGCCACTGTGACCATTGACTC. WT PCR product size: WT: 219 bp. WT and flox alleles must be analyzed in separate PCR reactions. ***Amelx**$^{flox}$*: Forward1: CTACAAGTGGACCTAAACCC; Forward2: GCAAGATCTTATATCCCCAAG; Reverse1: GAAGTTATGCTGCAGGTTAAC; Reverse2: CGTTACAGGAAGATCTCCTAG. PCR product size: WT: 174 bp, flox: 344 bp. WT and flox alleles can be analyzed in single PCR reaction. ***Amtn**$^{flox}$*: Catch-all PCR: Forward: CTACTGACACATAAATCTCATTG; Reverse: GTAGAAAAATCAGAACCATGATTT. PCR product size: WT: 1023 bp, flox: 1091 bp, Null: 210 bp. Flox PCR: Forward: GCACCGAGTAAAGTGGAGAAG; Reverse: ATGGCTTTATAACTTCGTATAATG. Flox

PCR product size: 520 bp. WT PCR: Forward: GTAAGGTTTGGAGAAATCATAT; Reverse: CAATCAGGACTAAACTGTCTGTG. WT PCR product size: 305 bp. WT and flox alleles must be analyzed in separate PCR reactions. ***Mmp20**$^{flox}$*: Catch-all PCR: Forward: CAATGGCGTACCAAGCCAGGAAA; Reverse: GAGTGCCTATAATAGCCAAAAC. PCR product size: WT: 1232 bp, flox: 1300 bp. Flox PCR: Forward: CTCTCGAGATAACTTCGTATAGC; Reverse: CTCTTTGAATTGGGAAAGACGC. Flox PCR product size: 495 bp. WT PCR: Forward: ATCCACTTCACCCATTACAAGTC; Reverse: GTATGCCTGTTGTTCCTCAG. WT PCR product size: 189 bp. WT and flox alleles must be analyzed in separate PCR reactions. ***Klk4**$^{flox}$*: Flox PCR: Forward: GAGAGACATAACTTCGTATAGC; Reverse: CACCCAGCCACAGCCCATTC. Flox PCR product size: 459 bp. WT PCR: Forward: CTAAGTGAAGATCTATGCCTC; Reverse: AGGGGGTTCGTGCAGTGACC. WT PCR product size: 260 bp. WT and flox alleles must be analyzed in separate PCR reactions.

**Recombinase activity induction**

Mice were given an intraperitoneal injection of 2.5 mg/25 g of body weight of tamoxifen (Sigma-Aldrich, T5648-5G) dissolved in corn oil (Sigma-Aldrich, C8267) at a concentration of 25 mg/ml.

**Antibodies**

For flow cytometry of mouse incisors, the rat monoclonal anti-EpCAM-APC conjugate (BioLegend, G8.8) at dilution 1:200 was used. For immunofluorescence the anti-EpCAM was used (eBioscience, # ab223582) at dilution 1:100. Nuclei were counterstained with DAPI (stock concentration 5 mg/ml, Sigma-Aldrich, D9542; dilution 1:1,000 or 10,000).

**Tissue thin sections, immunostaining and imaging**

For fluorescence and immunofluorescence imaging, mice were euthanized with $CO_2$ and perfused intracardially with ice-cold PBS followed by 4% PFA prepared in PBS. Hemimandibles were then dissected out and fixed with 4% paraformaldehyde (PFA) in PBS overnight at 4ºC before being decalcified at room temperature using 0.5 M EDTA for 2 weeks, embedded in paraffin or OCT and sectioned (7 µm).

For immunofluorescence, paraffin sections were rehydrated, and antigen retrieval was performed by sub-boiling slides in a microwave for 15 min in a citrate buffer (pH 6.2) containing 10 mM citric acid, 2 mM EDTA and 0.05% Tween-20. OCT sections were defrosted, washed, and no antigen retrieval was performed. Samples were blocked in 1X animal-free blocker (Vector Laboratories, SP-5030) supplemented with 2.5% heat-inactivated goat serum, 0.02% SDS and 0.1% Triton X-100. Primary antibodies were diluted in the same blocking solution without serum and sections were incubated at 4°C for 12 to 16 hours. Appropriate secondary antibodies from Fisher Scientific were used at 1:150 dilution. Nuclei were counterstained with DAPI. Cryosections and paraffin sections were mounted with ProLong Gold Antifade (Thermo Fisher Scientific, #P36930), covered and imaged. Paraffin and OCT sections were imaged using Leica-TCS SP5 confocal microscope, Zeiss LSM900, or Leica Dmi8, and analyzed using LasX or Fiji ImageJ.

**Fixed whole-mount tissue clearing, immunostaining and imaging**

Following fixation in 4% PFA, samples were washed with PBS pH 7.4 and incubated in CUBIC-L (TCI, #T3740) for 24 hours at 37°C for delipidation and decoloring. Next, the samples were washed with PBS pH 7.4 three times for 1 hour, followed by decalcification using CUBIC-B (TCI, #T3780) for four to five days at 37°C. After decalcification the samples were washed using PBS pH 7.4 three times for 1 hour, followed by a second round of delipidation using CUBIC-L for 24 hours at 37°C. After the second round delipidation, samples were washed using PBS pH 7.4 three times for 1 hour followed by immunostaining. The samples were first incubated with blocking solution composed of 1X Animal-free Blocker (Vector Labs, #SP-5030), 10% Normal donkey serum (Jackson, #017-000-121), 0.3% Triton X-100 (Thermo-Fisher, #A16046.AP), and PBS pH 7.4, for 24 hours at 4°C. Next, samples were incubated with antibodies diluted in antibody dilution buffer composed of 1X Animal-free Blocker, 0.3% Triton X-100, and PBS pH 7.4, for 24 hours at 4°C. Samples were washed with PBS pH 7.4 three times for 1 hour to wash out unbound antibodies. Finally, samples were incubated with secondary antibodies diluted in antibody dilution buffer for 24 hours at 4°C followed by three 1-hour long washes with PBS pH 7.4. Prior imaging, samples were cleared using CUBIC-R+ (TCI, T3983) until samples were fully cleared (roughly 2 hours), then mounted on a dish (Ibidi, 81156). Fixed whole mount samples were imaged with an LSM-900 with Airyscan using the super resolution settings. After image acquisition, the files were processed using the Airyscan processing option of the Zeiss Zen Imaging Software (Zeiss) and equally adjusted if needed using Fiji ImageJ.

**Incisor explant live-imaging**

Hemimandibles were collected and dissected in PBS containing 50% rat serum (Valley Biomedical, #AS3061SC, Special Processed) at 37°C. The cervical loop was separated from the rest of the mandible and embedded in 1.5% low-melting agarose (NuSieve, #50080) with 50% rat serum in a 1:1 ratio, overlaid with complete culturing medium and incubated at 37°C and 5% $CO_2$ for one hour. Complete culturing medium was prepared by mixing 50% DMEM/F12, 50% rat serum, 1% Glucose (Sigma Aldrich, # G8769), 500 µg/ml Ascorbic acid (Sigma Aldrich, #A4403), 1% Penicillin-Streptomycin (Sigma Aldrich, #516106), 1X Glutamax (ThermoFisher, # 35050061), and 1X NEAA (ThermoFisher, # 11140050). During imaging, sample was maintained in 5% $CO_2$, the temperature was maintained at 37°C using a heating plate (Bioptechs), and fresh medium was recirculated using Delta T pump (Bioptechs). Time-lapse imaging of explants was carried out using a Nikon A1R upright microscope. Images were recorded every 15 minutes for 5 hours, using a 25x water-immersion objective (NA = 1.1), excitation wavelength of 880 nm and emission wavelength of 440 nm. Data were processed and videos were generated in ImageJ.

***In vitro* explant cultures**

Hemimandibles were collected in PBS and cervical loops were dissected and cultured for 24 hours at 37°C and 5% $CO_2$ in DMEM/F12 (Gibco, 21041025) supplemented with 5% heat inactivated FBS and 1µM 4-hydroxy-tamoxifen to induce Cre recombination. After an additional 24 hours of incubation, the cervical loops were fixed with 4% PFA in PBS (pH 7.4) for 24 hours at 4°C. Fixed samples were processed for antibody staining as detailed in the sample processing for whole-mount tissue clearing and immunostaining.

**Flow cytometry analysis and cell sorting**

Sample preparation for flow cytometry was described earlier (Sharir et al., 2019). In short, the bulbous portion, as well as the lateral wing-shaped epithelium, and the surrounding mesenchyme were dissected from the incisor, collected in cold $Ca^{2+}$- and $Mg^{2+}$-free HBSS (pH 8.0, UCSF Cell Culture Facility, CCFAJ005-16CT01) and digested enzymatically at 37 °C in collagenase P (2 mg/ml in PBS; Sigma Millipore, 11213857001) for 30 min, followed by collagenase inactivation with 2% FBS. The digested tissue was mechanically disrupted using a 1 ml pipette tip and centrifuged at 400g for 3 min. Cells were resuspended in HBSS (pH 8.0) containing 2% FBS. Cell suspensions were stained with rat monoclonal anti-EpCAM (BioLegend; clone G8.8; AF647 conjugate) for 30 min on ice followed by centrifugation at 400g for 3 min. Cells were filtered through a 40 µm mesh (BD Falcon, 22363547) and stained with DAPI (1:10,000) prior flow cytometric analysis and sorting (BD FACSAria2 SORP).

**Quantitative PCR**

Total RNA from 1,000 – 5,000 sorted cells was extracted using RNeasy Mini Kit (Qiagen, #74104) according to the manufacturer's protocol. cDNA was synthesized with High-capacity cDNA Reverse Transcription Kit (Applied Biosystems, #4368814). qPCR reactions were performed using iTaq Universal SYBR Green Super-mix (Bio Rad, #1725121) in 384-well plates on a QuantStudio 6 Flex Real-Time PCR System (Thermo Fisher Scientific). Primers used for qPCR in 5'-3' orientation: *Gapdh*: AATGGTGAAGGTCGGTGTG and GTGGAGTCATACTGGAACATGTAG; *Ambn*: GAGAAGTCCGTGCAACCATA and TGAGCCTTGAGACAATGAGAC; *Klk4*: ACACTTTCCTTGTCCCATAGAC and CTGTGTACCACCTCAGTATGTTC.

**Synchrotron Micro-computed tomography (SMCT)**

Mice at least eight weeks old were anesthetized with $CO_2$ and perfused intracardially with phosphate-buffered saline (PBS) followed by 4% paraformaldehyde (PFA) in PBS. Jaws were collected and soft tissue was removed manually and stored in 70% ethanol (VWR, Radnor, PA) at room temperature. Hemimandibles were positioned with a small length of aluminum wire (approx. 1 cm, >99.99%, Sigma-Aldrich) on custom-designed, radiotransparent 3D printed holders and held in place with parafilm. The holders were wedged with Styrofoam packing peanuts into 2 mL Eppendorf tubes that were previously filled with ~0.5 mL ethanol to maintain a saturated atmosphere and prevent the samples from drying out. Samples were transported and stored at the beamline at room temperature. SMCT datasets were collected at Argonne National Laboratory, Advanced Photon Source, beamline 2BM-A,B (Nikitin et al., 2022). Experiments were performed with a monochromatic beam (25.51 keV). A 2x objective lens was used to achieve an isotropic voxel size of 1.73 μm. Radiographs were captured with a Flir camera (3.45 μm pixels) using a frame time of 0.20 s per projection, with 1500 projections over 180˚ (0.120˚ increment between frames). Dark field and flat-field images were collected at the beginning of each scan to normalize projection intensity. The rotation axis was approximately parallel to the long axis of the hemimandible, with the incisal tip pointing down. Collection lasted approximately 7 minutes per scan, and three scans stacked vertically with ~200 μm overlap between frames were collected for each hemimandible to capture secretory, transition, and maturation stages. Tomographic reconstruction was performed using a fourierrec algorithm within the open-source Python package tomocupy (Nikitin, 2022). Reconstruction included a stripe removal step, using the Fourier wavelet method. Reconstructions were stored as 32-bit float TIF image sequences.

**Scanning electron microscopy**

Heads from mice at least eight weeks old were collected and shipped to Northwestern University on dry ice. Heads were stored at -80˚C and thawed at room temperature for dissection. Soft tissue was removed manually from the hemimandibles, which were then dehydrated in an ethanol series (10%, 30%, 50%, 70%, 90%, 100% (v/v) ethanol in deionized water, at least 15 min. per step) before being infiltrated with LR white resin (2:1 and 1:1 mixtures ethanol: LR white mixtures, at least 12 hours at room temperature with gentle rocking). The hemimandibles were then embedded in 100% LR white in gelatin capsules and left to cure overnight at 65˚C. The gelatin was removed, and the LR white pellet was embedded in epoxy resin at room temperature overnight. Coronal sections (normal to the long axis of the jaw) were cut with an Isomet saw (Buehler) diamond wafering blade (Allied High-Tech Products) at the point of incisor eruption. Sections were ground with SiC papers (600, 800, 1000 grit) and polished (3 μm, 1 μm aqueous diamond suspensions, 0.05 μm aqueous silica suspension) before being etched with 10 μL of 250 mM lactic acid (pH ~3.7) for 5 seconds. Samples were stored under vacuum before being coated with 5 nm Au/Pd using a Denton Desk IV sputter coater (Denton Vacuum). SEM was performed using a Hitachi S4800-II (Hitachi High-Tech) scanning electron microscope. Images were captured with an accelerating voltage of 5 kV, a working distance of 1.5-4 mm, and an emission current of ~10 μA using secondary electron contrast.

**Statistical analysis and compliance with the ARRIVE guidelines 2.0**

Study design and sample size: For fluorescence imaging we used at least two males or females from two independent F2 litters (N=4). F1 generation contains stably inserted mutant allele, and this one was genotyped and sequenced. For all other experiments we used at least three males or females from F2 litter (N=3). Inclusion and exclusion criteria: Wildtype mice or mice negative for the specific allele determined by genotyping/sequencing were used as controls. Only offspring of mutant founders that contained the entire exogenous DNA inserted correctly (confirmed by genotyping/sequencing), was propagated and included in the study. Randomization and blinding: Prior fluorescence microscopy, SEM and SMCT analyses, samples were randomized by hiding the genotype and replacing it with number. Blinding was done by Tomas Wald. No criteria for excluding animals from quantitative analyses were set and no animals were excluded from qPCR analyses. Statistical analysis was determined in in GraphPad Prism 9 (GraphPad Software, La Jolla, CA). The value N represents the number of animals or preparations. Normally distributed data were analyzed using parametric Student's t-test with Welch's correction or one-way ANOVA with Tukey's multiple comparisons test. The non-parametric Mann–Whitney U-test was used if the data did not fit a normal distribution. Significance was taken as $P < 0.05$ with a confidence interval of 95%. Data are presented as mean ± SD for parametric data or as median ± interquartile range for non-parametric data.
